# Serum extracellular vesicle proteomics reveals treatment-associated biomarkers in pancreatic ductal adenocarcinoma

**DOI:** 10.64898/2026.11.06.764966

**Authors:** Sehong Min, Komal Abhange, Tai-Tu Lin, Jianliang Dai, Isaac K. Attah, Anders Bogard, Vaibhav Sahai, Kyle C. Cuneo, Suyu Liu, David M. Lubman, Tujin Shi

**Author notes:** **Corresponding authors:** Dr. David M Lubman Department of Surgery, University of Michigan Medical Center Ann Arbor, MI 48109, **Email:**; Dr. Tujin Shi, Department of Chemistry and Chemical Biology Indiana University, Indianapolis, IN 46202, **Email:**.

## Abstract

**Background:** Longitudinal biomarkers of treatment response in pancreatic ductal adenocarcinoma (PDAC) are needed. Extracellular vesicle (EV)-enriched serum proteomics may provide a minimally invasive approach for monitoring systemic and tumor-associated responses to therapy.

**Methods:** Paired pre-treatment and post-treatment serum samples were analyzed from 51 patients with pancreatic cancer, including 25 treated with FOLFIRINOX and 20 treated with gemcitabine-based therapy. EVs were enriched using the Mag-Net method and profiled by data independent acquisition mass spectrometry. After batch correction and filtering, 941 proteins were retained for differential and pathway analyses. Paired linear models and gene set enrichment analyses were used to assess treatment-associated changes. Baseline protein abundance and longitudinal changes were evaluated in relation to overall survival (OS) using Cox models.

**Results:** A total of 2,089 EV-associated proteins were identified, with an average of 1,186 proteins quantified per sample. Across the cohort, 26 proteins differed between post- and pretreatment samples. Proteins that increased after treatment were primarily related to extracellular matrix organization and cell adhesion, whereas proteins that decreased after treatment were associated with actin cytoskeletal organization, coagulation, and hemostasis. Treatment-associated changes were more pronounced in the FOLFIRINOX group, in which 32 proteins were differentially expressed. FOLFIRINOX was characterized by coordinated reductions in platelet- and coagulation-associated proteins, including PF4, PF4V1, PPBP, GP5, ITGA2B, and ITGB3, accompanied by enrichment of platelet activation, coagulation, and hemostasis pathways among down-regulated processes. No individual proteins met the same significance threshold in the gemcitabine group. A composite seven-protein platelet score decreased significantly after FOLFIRINOX but not for gemcitabine, and the between-regimen difference in score change was significant. Higher pretreatment ITGB1 abundance was associated with longer OS after multiple-testing correction (HR, 0.112; BH-FDR = 0.0063). In multivariable analysis, greater treatment-associated platelet-score depletion was independently associated with shorter OS (HR per unit increase in Δscore, 0.56; P = 0.030).

**Conclusions:** Longitudinal serum EV-enriched proteomics identified treatment-associated biological changes in PDAC, including a coordinated platelet-related response predominantly observed after FOLFIRINOX. Baseline ITGB1 abundance and longitudinal platelet-score changes represent potentially complementary prognostic and pharmacodynamic markers. These findings are exploratory and require validation in larger, independent cohorts incorporating platelet counts, coagulation measurements, and targeted proteomic assays.

## Introduction

Pancreatic cancer is currently the third leading cause of cancer related to death in the US [1]. A significant portion of these patients present with unresectable locally advanced disease where the standard of care includes a combination of chemotherapy and radiation therapy. A major challenge in the management of pancreatic cancer is the early assessment of treatment response. Novel markers of early treatment response are critically needed to make better informed decisions regarding the type and sequencing of therapies, given the high toxicity and variable effectiveness associated with these treatments. Computed tomography (CT) is currently used to stage and reassess patients following treatment. However, current imaging modalities are not effective at identifying early, small-volume metastatic disease [2–6]. Novel biomarkers of early treatment response are greatly needed to improve the management of patients with pancreatic cancer. Early identification of treatment response based on predictive markers would help to select the most appropriate therapy and potentially limit toxicity in non-responding patients.

Different types of membrane nanovesicles are secreted by most types of cells, including both normal and cancer cell-derived nanovesicles into the bloodstream. There are different categories of vesicles where in the present work we are focused on extracellular vesicles (EVs) [7–9]. EVs contain a variety of distinctive cargo including proteins. Thus, tumor-derived serum proteins in EVs can be used to provide a minimally invasive method for detection, prognosis or therapeutic monitoring [7–15]. EVs are selectively loaded with cargo related to the disease state which significantly reduces the complexity compared to searching for markers directly in serum and they can be obtained as part of routine blood draws. For pancreatic cancer and most other internal organs, monitoring EVs from serum is appropriate [11]. However, for the isolation of EVs from serum the presence of high abundance blood proteins presents a special challenge that can interfere with analysis.

Tumor-derived EVs can be isolated from patient serum using a number of different techniques including ultrafiltration, ultracentrifugation (UC) [8, 11], sucrose gradient fractionation, size exclusion chromatography (SEC) [16, 17], antibody-based extraction [18, 19], asymmetric field flow [20], polymeric precipitation [21], and various microchip devices [22] among others [23–25]. Although there are many EV isolation methods available there are still significant problems for clinical use. Most EV isolation methods require a significant volume of sample and time to perform the isolation and are labor-intensive. In addition, they may lack robustness and reproducibility, which is an important technical challenge for their broad application in clinical use. In our previous studies we have used multiple cycles of ultracentrifugation to provide the required purification for analysis of EVs in mass spectrometry discovery-based work [26]. However, this method is time-consuming and labor-intensive and requires at least 0.5 mL of serum and results in a low yield of vesicles.

More recently, several bead-based isolation methods have been developed for isolation of EVs. Of note is the use of Mag-Net, which uses strong anion exchange (SAX) beads, functionalized with quaternary ammonium to isolate EVs through electrostatic interactions while depleting blood proteins [24]. This approach is particularly effective at reducing the lipoprotein content where these proteins and their complexes are difficult to eliminate based on size-based methods. By enriching EVs, Mag-Net enables the detection of proteins that would otherwise fall below the dynamic range of liquid chromatography–tandem mass spectrometry (LC–MS/MS) analysis of unfractionated serum. The method is relatively simple and provides robust, reproducible EV isolation, followed by on-bead lysis and digestion of EV proteins for mass spectrometry analysis.

An important goal of our current research is to use information from EVs to personalize therapy for pancreatic cancer. There are several systemic therapy options for patients with pancreatic cancer including gemcitabine nab-paclitaxel and FOLFIRINOX. These are the regimens recommended in the NCCN guidelines and are considered effective therapies for this disease [27, 28]. In addition, there are dozens of novel therapies currently in clinical trials. In the patients included in our prior EV work, overall survival (OS) approached 2 years, which is higher than previously reported in locally advanced patients [29]. As more effective therapies are developed, the ability to personalize and adapt treatment for an individual patient will be vital to limit toxicity and maximize effectiveness.

In this study, we used serum EV proteomics to investigate treatment-associated changes in patients with pancreatic ductal adenocarcinoma (PDAC). EVs were isolated from only 50 μL of serum using the Mag-Net method, and their protein content was analyzed by data independent acquisition (DIA)-MS. More than 2,000 EV proteins were identified across the PDAC serum samples, with 26 proteins showing significant differences between post- and pretreatment samples. The main findings involved extracellular matrix remodeling and, particularly after FOLFIRINOX treatment, a coordinated reduction in platelet- and coagulation-associated proteins. Baseline ITGB1 abundance and an exploratory multimarker score were associated with OS. These findings suggest that serum EV proteomics can capture complementary biological information, including pretreatment disease-associated risk and treatment-related pharmacodynamic responses. They further support the potential use of EV protein biomarkers for monitoring therapeutic responses in PDAC.

## Methods

### Reagents and Chemicals

All chemicals were purchased from Sigma-Aldrich (St. Louis, MO) unless otherwise stated. Mass spectrometry-grade trypsin (Cat. V5280) was purchased from Promega Corporation (Madison, WI).

### Study Design and Serum Sample Collection

The serum samples were obtained from patients with PDAC after providing informed consent on a University of Michigan Medicine Institutional Review Board study (HUM00025339). The samples were collected in serum tubes, processed using a standardized workflow within 2 hours of collection, stored at −80 °C, and used without freeze–thaw cycles.

Paired pre- and post-treatment serum from 51 patients with PDAC were analyzed. EVs were enriched from 50 µL of serum using the Mag-Net protocol, automated on a KingFisher Flex system. Peptides were analyzed by data-independent acquisition (DIA) on a Vanquish Neo nanoLC system coupled to an Astral mass spectrometer, and raw data were processed with DIA-NN.

### EV Enrichment

EV enrichment was performed using the Mag-Net protocol [30], a magnetic bead-based method that enables charge- and size-based EV enrichment via an automated workflow. Briefly, serum samples were centrifuged at 2,000 *g* for 30 min. 50 µL of serum supernatant was mixed 1:1 (v/v) with binding buffer (100 mM Bis-Tris Propane, pH 6.3, 150 mM NaCl). This process was performed in two cycles on a Thermo Fisher Scientific KingFisher (KF) Flex according to the manufacturer’s protocol with minor modifications as follows. KF Cycle 1: 12.5 µL of MagReSyn SAX beads (ReSyn Biosciences, MR-SAX002) were first equilibrated twice in 500 µL of equilibration/wash buffer (WB, 50 mM Bis-Tris Propane, pH 6.5, 150 mM NaCl), then incubated with the serum-binding buffer mix for 45 minutes at room temperature. The beads were washed with WB of 500 µL for 5 min three times. The captured membrane particles on the beads were lysed and reduced in 100 µL of lysis buffer (50 mM Tris, pH 8.5, 1% SDS, 10 mM Tris (2-carboxyethyl) phosphine (TCEP)) for 1 h at 37 °C. After KF Cycle 1, the samples were removed from KF, alkylated with 15 mM iodoacetamide (IAA) in the dark for 30 min under gentle agitation, and mixed with 70% acetonitrile (ACN) before being returned to the KingFisher system for Cycle 2. KF Cycle 2: The bead-protein-ACN mixture was incubated for 10 min at room temperature to precipitate proteins onto the bead surface. The beads were then washed three times with 500 µL of 95% ACN and twice with 500 µL of 70% ethanol. Following the washes, the bead-bound proteins were digested for 1 h at 47 °C in 100 mM ammonium bicarbonate with 3.75 µg of trypsin. Digestion was quenched by adding trifluoroacetic acid to a final concentration of 0.5%, and peptides were then stored at −80 °C until LC-MS/MS analysis.

EV enrichment was performed in two experimental batches: batch 1 (21 patient pairs, 42 samples) and batch 2 (30 patient pairs, 60 samples, in addition to 3 pairs from batch 1 for batch control). Critically, both pre- and post-treatment samples from the same patient were processed within the same batch to preserve the paired structure and prevent batch effects from confounding within-patient comparisons.

### LC-MS/MS Analysis

Peptides were analyzed by LC-MS/MS on an Orbitrap Astral mass spectrometer (Thermo Fisher Scientific) coupled to a Vanquish Neo liquid chromatography system (Thermo Fisher Scientific). 500 ng of peptides per sample were separated on a PepMap ES906 analytical column for reverse-phase elution using a 50 SPD separation method, with each sample analyzed in a 23-minute active gradient. The Vanquish Neo was operated in the trap-and-elute mode with a PepMap Neo Trap Cartridge (Thermo Fisher Scientific), which was used for sample trapping, and operated in the “backwards flush” mode. The two mobile phases used were (A) 0.1% formic acid in water and (B) 0.1% formic acid in acetonitrile. The separation gradient profile used as follows: (min, Flow Rate, %B): 0.0, 3.50, 1.0; 0.001, 2.8, 1.0; 1.001, 1.30, 4.0; 2., 0.80, 8.0; 3.0, 0.80, 8.0; 18.001, 0.80, 22.5; 22.001, 0.80, 35.0; 23.001, 1.10, 45.0. Column Wash; 23.501, 2.80, 99.0; 24.701, 2.80; 99.0; 25.701, 3.50, 1.0.

Data were acquired in DIA mode with the following parameters. An EASY-Spray ion source (Thermo Fisher Scientific) was used with the spray voltage at 2.2 kV. The Ion transfer tube temperature was set to 300 °C. Full scan spectra were acquired using the Orbitrap analyzer over a scan range of 380 to 980 m/z at 240,000 resolution. A normalized AGC target of 500% was used, with the maximum injection time set to 5 ms and the RF lens was set at 45%. For the nDIA settings, the DIA window type was set to “Auto”, window placement optimization at “On”, window overlap at 0, and the isolation window at 2 m/z. HCD fragmentation was performed at a normalized collision energy of 25%, with the scan range for the Astral mass analyzer set at 150 – 2000 m/z. Loop control was set at 0.6 seconds. To minimize potential batch effects and analytical biases, the order of sample runs was randomized across the LC-MS/MS experiments.

### Data Processing

Raw data were processed and searched against the UniProt human proteome database (FASTA file dated 2023-09-01; 20,441 entries) and common contaminants using DIA-NN (v2.0) in library-free mode [31]. Each batch was searched as a separate patch. In silico digestion was performed with trypsin specificity (cleavage after K and R), allowing up to two missed cleavages, with peptide lengths of 7–30 amino acids and precursor m/z range of 380–980 (charge states 2–4). Fixed modification includes cysteine carbamidomethylation (+57.021 Da). Variable modifications included methionine oxidation (+15.995 Da) and protein N-terminal acetylation (+42.011 Da), with a maximum of three variable modifications per peptide. Precursor-level false discovery rate was controlled at 1%. Protein inference was performed at the gene level using heuristic protein inference. Quantification was performed using the Quant UMS algorithm in high-precision mode. Match-between-runs was enabled. MS1 and MS2 mass accuracies and scan window were determined automatically by DIA-NN. RT-dependent cross-run normalization was applied within DIA-NN.

Protein intensities were log₂-transformed for downstream analysis. To account for technical variation between the two EV enrichment batches, we applied a median-based batch correction approach. Empirical Bayes batch correction (ComBat) was initially evaluated but failed to converge across multiple detection thresholds, likely due to the relatively small number of samples per batch and sparse missing value patterns. We therefore implemented a manual median-alignment strategy: for each protein detected in at least 70% of samples, the batch 2 median was shifted to align with the batch 1 median, using batch 1 as the reference. This approach preserved within-patient paired differences (since paired samples resided in the same batch) while correcting systematic inter-batch offsets. Following batch correction, global median normalization was applied across all samples to equalize the median log₂ intensity across samples and ensure comparability.

Abundant serum contaminant proteins that were co-isolated during EV enrichment were identified and removed prior to differential expression and downstream analyses. These included hemoglobin subunits (HBA1, HBA2, HBB, HBD, and related chains), albumin (ALB), and apolipoproteins (APOA1, APOA2, APOB, APOC family members, APOE, and others), a total of 17 proteins.

### Data Analysis

#### Differential Expression Analysis

For within-treatment paired analysis, treatment-induced proteomic changes were assessed separately for each regimen using the limma package (v3.58.1) in R. For each treatment group, a paired linear model was fit with patient as a blocking factor: Expression ∼ Patient + SampleType, where SampleType encoded pre- vs. post-treatment status. Moderated t-statistics were computed using empirical Bayes shrinkage. Differentially expressed proteins (DEPs) were defined by a dual threshold of Benjamini–Hochberg adjusted p-value (FDR) < 0.05 and absolute log₂ fold change > log₂(1.3) (corresponding to a ±30% change on the linear scale).

To compare treatment-induced changes between FOLFIRINOX and gemcitabine, a delta matrix was computed for each protein as Δ = post-treatment intensity − pre-treatment intensity, yielding one value per patient. A linear model was then fit on the delta matrix: Δ ∼ Batch + Treatment, where Batch accounted for the EV enrichment batch and Treatment encoded FOLFIRINOX vs. gemcitabine. This model was extended to include age as a covariate (Δ ∼ Batch + Age + Treatment), given a significant age imbalance between treatment groups (median 66 vs. 74 years; Wilcoxon p = 0.003).

#### Pathway Enrichment Analysis

Over-representation analysis (ORA) and gene set enrichment analysis (GSEA) were performed using functions from the clusterProfiler package. The moderated t-statistic from limma was used as the ranking metric for GSEA. GSEA was additionally performed on the Cox proportional hazards regression results (*see Survival Analysis*), using a composite ranking metric of sign(coefficient) × −log₁₀(p-value) from the univariate Cox screening.

### Statistical Analysis

Descriptive statistics were used to summarize patient characteristics for the cohort. The Wilcoxon rank-sum test was used to compare continuous variables, while Fisher’s exact test or the chi-square test was used to compare categorical variables between groups. As the markers were measured before and after treatment, a paired linear model was used to assess changes in marker levels over the course of treatment. Benjamini–Hochberg method was used to adjust the p-value for multiple testing. The Kaplan–Meier method was used to estimate survival functions, and the log-rank test was used to compare survival between groups. Cox regression model was used to assess the association between markers as well as patient characteristics with survival.

#### Platelet EV Signature Score

A composite platelet EV signature score was defined as the mean z-score of seven platelet-associated proteins that were significantly downregulated after FOLFIRINOX treatment: PF4, PF4V1, ITGA2B, ITGB3, GP5, PPBP, and SPARC. For each protein, z-scores were computed across all 90 samples (mean = 0, SD = 1), and the signature score for each sample was the arithmetic mean of these seven z-scores. The treatment-induced change in platelet signature (ΔPlatelet score) was calculated as the post-treatment score minus the pre-treatment score for each patient. Within-treatment pre- vs. post-treatment differences were assessed by paired Wilcoxon signed-rank tests, and between-treatment differences in ΔPlatelet score were assessed by the unpaired Wilcoxon rank-sum test.

#### Survival Analysis

OS was analyzed in the full cohort of 45 patients (36 events). Survival time was measured in months from diagnosis to death or censoring at last follow-up.

#### Univariate Cox Screening

Univariate Cox proportional hazards regression was performed for each of the proteins using pre-treatment expression levels, adjusted for age and treatment regimen. Proteins with nominal p < 0.05 were considered prognostic candidates. Pathway enrichment analysis (ORA and GSEA) was then applied to these prognostic hits to characterize the biological pathways associated with survival.

#### Kaplan–Meier Analysis

Patients were dichotomized at the median ΔPlatelet score into “large decrease” and “small decrease” groups for Kaplan–Meier visualization. Survival curves were compared using the log-rank test. Kaplan–Meier analyses were also performed by treatment regimen.

All statistical analyses were performed in R (v4.3.3). Key packages included limma (v3.58.1) for differential expression, clusterProfiler (v4.10.1) and org.Hs.eg.db. Two-sided tests were used throughout unless otherwise specified. A p-value < 0.05 was considered statistically significant. FDR < 0.05 is used to identify the predictive markers accounting for multiple testing.

## Results

### Pancreatic cancer cohort characteristics and serum EV proteome profiling

Paired pre- and post-treatment serum samples were collected from 51 patients with pancreatic cancer, comprising 25 who received FOLFIRINOX, 20 gemcitabine-based, and 6 other regimens. The cohort was predominantly white non-Hispanic or Latino (49/51, 96%). Patients treated with FOLFIRINOX were more often male (16/25, 64%) than those treated with gemcitabine (11/20, 55%). Patients receiving gemcitabine-based chemotherapy were older than those receiving FOLFIRINOX (median age at collection 74 vs. 66 years; P = 0.003), consistent with the clinical practice of reserving the more intensive FOLFIRINOX regimen for younger patients [32]. At the time of analysis, 18/25 (72%) of FOLFIRINOX-treated and 18/20 (90%) of gemcitabine-based treated patients were deceased. EV enrichment was performed across two batches (20 and 31 pairs each), with pre- and post-treatment samples from the same patient processed within the same batch.

MS-based proteomic profiling identified 2089 serum EV proteins across all 51 paired pancreatic cancer patient samples using the Mag-Net method, with an average of 1186 proteins quantified per sample and comparable coverage between the pre- and post-treatment timepoints. Three patient pairs were processed in both batches as bridge samples to enable assessment of batch effects. After retaining 958 proteins detected in at least 70% of samples within each batch, we applied a median-based batch correction followed by median normalization. Signal intensity distributions, the correlation of bridge samples across batches, and principal component analysis (PCA) before and after correction confirmed that batch structure was substantially reduced. After removal of abundant contaminant proteins (hemoglobin, albumin, and apolipoprotein species), 941 proteins were retained for differential and pathway analyses.

### Whole cohort characterization of the pre-versus post-treatment serum EV proteome

Canonical EV-positive markers (CD9, CD63, CD81, PDCD6IP (ALIX), SDCBP (Syntenin-1), HSPA8) were consistently detected across the majority of samples, supporting EV enrichment of the preparations (**Fig. 2A**). Among previously reported pancreatic cancer-associated markers, TFPI, TIMP1, LRG1, THBS2, GPC1, and S100A4 were detectable and are shown pre- vs post-treatment (**Fig. 2B**). Expression levels of both the EV and pancreatic cancer markers were comparable between pre- and post-treatment samples.

**Fig. 1.**
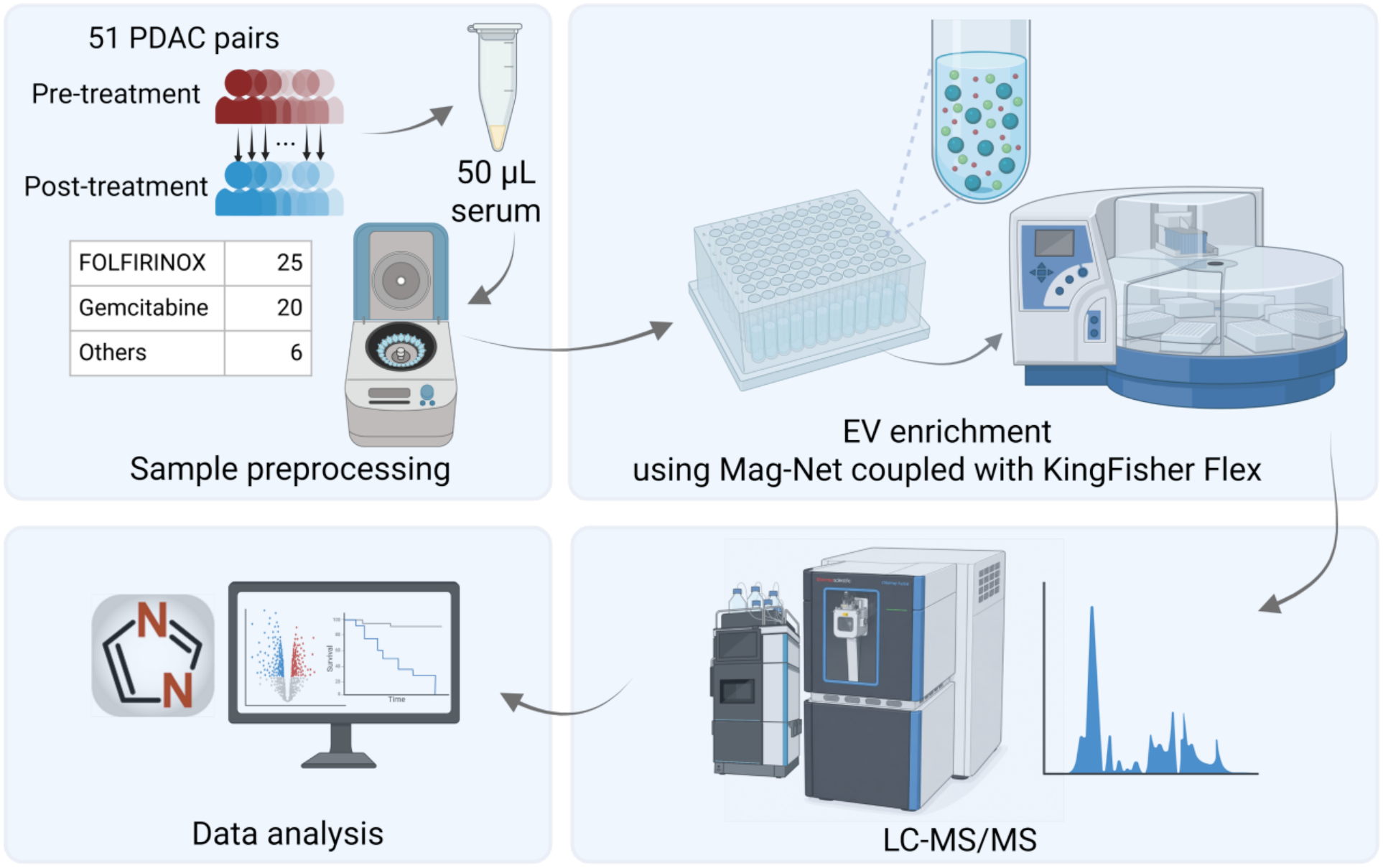
Schematic workflow for the study. Paired pre- and post-treatment serum from 51 pancreatic ductal adenocarcinoma (PDAC) were analyzed. Extracellular vesicles (EVs) were enriched from 50 µL of serum using the Mag-Net protocol, automated on a KingFisher Flex system. Peptides were analyzed by data-independent acquisition (DIA) on a Vanquish Neo nanoLC system coupled to an Astral mass spectrometer, and raw data were processed with DIA-NN.

**Fig. 2.**
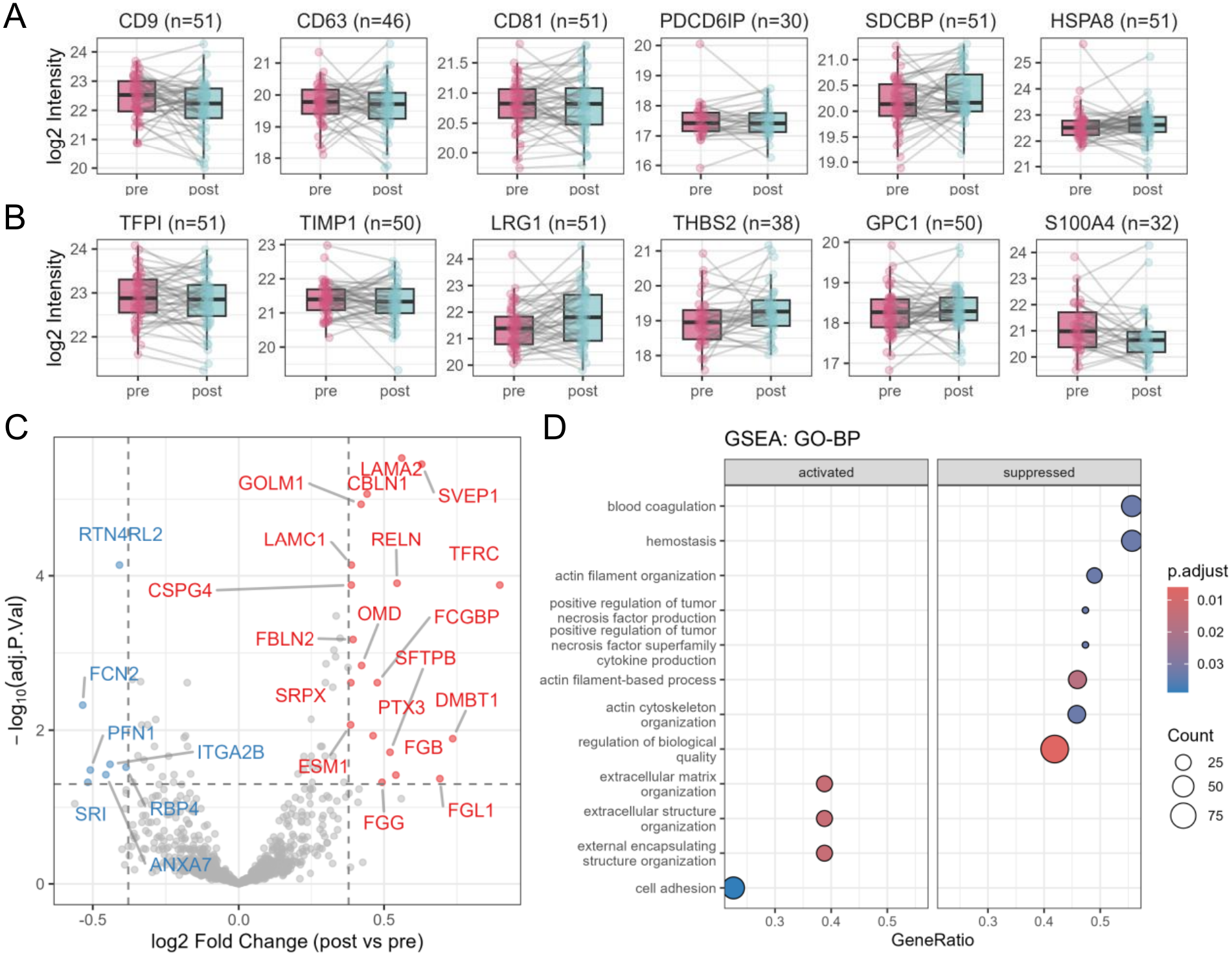
Whole cohort landscape of the pre- and post-treatment EV-enriched serum proteome. (A) EV-positive markers (CD9, CD63, CD81, PDCD6IP (ALIX), SDCBP (syntenin), HSPA8) in pre- vs post-treatment samples. (B) PDAC-associated markers (TFPI, TIMP1, LRG1, THBS2, GPC1, S100A4) in pre- vs post-treatment samples. (C) Volcano plot of the pooled paired post- vs pre-treatment comparison. Dashed lines denote significance thresholds (BH-FDR < 0.05 and |log2FC| > log2(1.3)), and 26 proteins reached significance. (D) GSEA (GO-BP) of the pooled comparison.

To characterize the overall treatment-associated change, we compared post- vs pre-treatment serum EV proteomes across the whole cohort using a paired linear model. A total of 26 proteins reached significance (BH-FDR < 0.05, |log2FC| > log2(1.3)) (**Fig. 2C**). The up-regulated proteins (19 proteins higher in post-treatment) were dominated by extracellular matrix and cell-adhesion components, including the laminins LAMA2 and LAMC1, FBLN2, SVEP1, and RELN, whereas the down-regulated proteins (7 proteins higher in pre-treatment) were enriched for actin cytoskeleton regulators, including PFN1, ANXA7, and SRI. Gene set enrichment analysis (GSEA, GO-BP) was consistent with differential analysis results, showing that extracellular matrix organization and cell adhesion were enriched among up-regulated proteins, whereas actin cytoskeleton organization, blood coagulation, and hemostasis were enriched among down-regulated proteins (**Fig. 2D**). Because treatment differed across the cohort, we next asked whether this signal was uniform across the two regimens.

### Treatment-associated EV proteome changes were more pronounced in the FOLFIRINOX group

To determine whether the whole cohort treatment-associated changes were uniform across regimens, we performed paired post- vs pre-treatment analyses separately within the FOLFIRINOX (25 pairs) and gemcitabine (20 pairs) groups. Because treatment assignment was not randomized, we first compared the two groups at baseline (i.e. pre-gemcitabine vs pre-FOLFIRINOX). Thirty proteins differed between pre-gemcitabine and pre-FOLFIRINOX samples (BH-FDR < 0.05, |log2FC| > log2(1.3)). The difference was not attributable to sample quality, detection coverage, normalization, batch, demographics, or EV fraction composition. These proteins were dominated by ubiquitin–proteasome and chaperone components and were unrelated to the platelet-associated proteins described below (r = 0.05). No proteins differed between the two groups after treatment. Because every comparison below is paired, with each patient serving as their own control, this baseline difference does not contribute to the within-regimen results. PCA of each treatment showed that pre- and post-treatment samples overlapped. EV and pancreatic cancer-associated markers were comparable between timepoints in both groups (**Fig. 3A-B**).

**Fig. 3.**
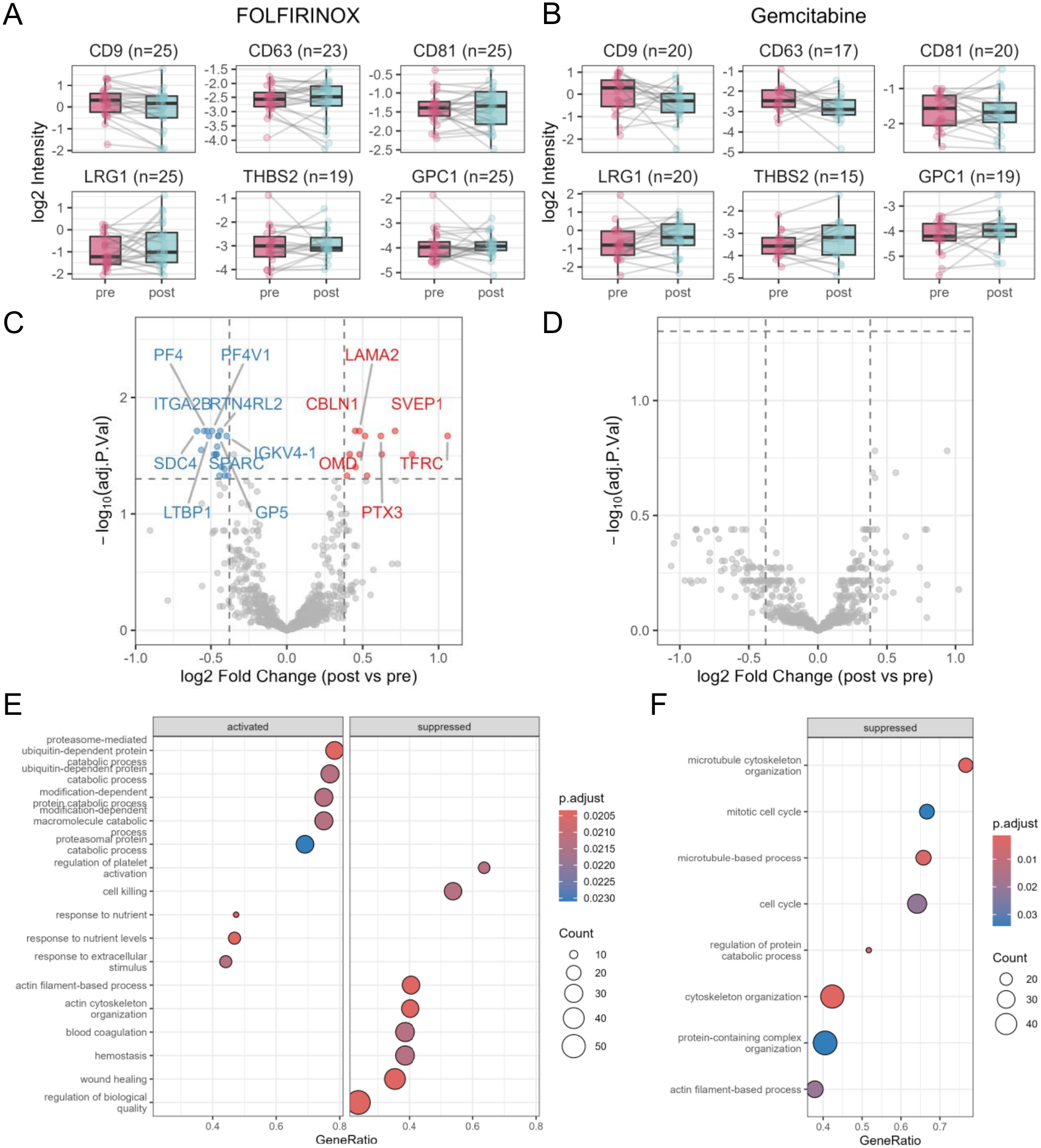
Paired analyses stratified by treatment reveal a FOLFIRINOX-specific platelet/coagulation response. (A-B) EV-positive markers and PDAC-associated markers in pre- vs post-treatment samples; FOLFIRINOX (A), gemcitabine (B). (C) Volcano plot of the paired post- vs pre-treatment comparison within FOLFIRINOX (n = 25 pairs); 32 proteins met significance (BH-FDR < 0.05, |log2FC| > log2(1.3)). (D) The corresponding volcano for gemcitabine (n = 20 pairs). (E) GSEA (GO-BP) of the FOLFIRINOX comparison. (F) GSEA (GO-BP) of the gemcitabine comparison.

The FOLFIRINOX comparison showed that 32 proteins were differentially abundant (BH-FDR < 0.05, |log2FC| > log2(1.3)) (**Fig. 3C**). Notably, platelet and coagulation-related proteins were decreased in the post-FOLFIRINOX samples, including the platelet α-granule chemokines PF4, PF4V1 and PPBP, and the platelet receptors and integrin subunits GP5, ITGA2B, and ITGB3. Consistently, GSEA (GO-BP) identified regulation of platelet activation, blood coagulation, and hemostasis among the most significantly down-regulated processes (**Fig. 3E**). Meanwhile, extracellular matrix and structural proteins, including COL1A1, LAMA2, SVEP1, HSPG2, and RELN, increased after treatment, consistent with the enriched extracellular matrix organization signal observed in the whole-cohort analysis. In contrast, the gemcitabine comparison did not show proteins meeting the same significance threshold (**Fig. 3D**). At the pathway level, GSEA identified down-regulation of cytoskeleton organization, microtubule-based processes, and cell-cycle-associated terms (**Fig. 3F**). None of the platelet- or coagulation-associated processes enriched in the FOLFIRINOX group were observed in the gemcitabine group. These results were consistently observed in overrepresentation analysis (ORA KEGG). Together, these analyses indicate that the platelet and coagulation signature of the serum EV response is predominantly associated with FOLFIRINOX, whereas the gemcitabine-associated changes involved cytoskeletal and cell-cycle processes.

### Baseline EV protein abundance is prognostic for OS

We next examined OS of FOLFIRINOX or gemcitabine recipients (time origin, start of therapy; 36 deaths among 45 patients). Treatment showed a marginally significant difference in OS, with longer survival under FOLFIRINOX (log-rank P = 0.022) (**Fig. 4A**). Among the covariates examined (treatment, age at diagnosis, and gender), only treatment reached a nominal raw P <0.05. Because FOLFIRINOX recipients were significantly younger than gemcitabine recipients (median 66 vs 74 years; P = 0.004), consistent with typical clinical practice, this treatment comparison is observational and cannot be interpreted causally.

**Fig. 4.**
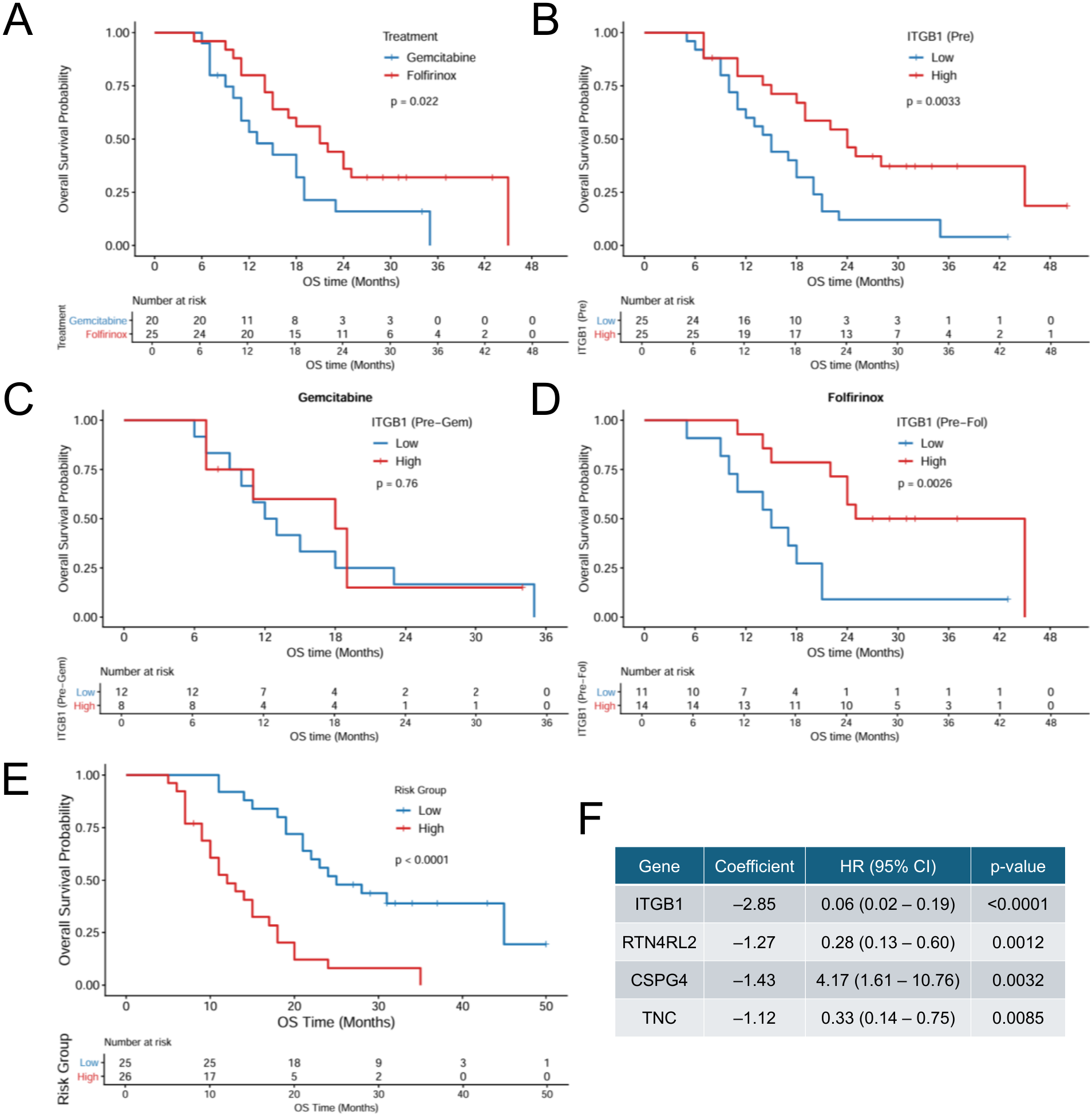
EV-enriched ITGB1 is associated with patient survival. (A) Kaplan-Meier overall survival (OS) by treatment group (FOLFIRINOX, n = 25; gemcitabine, n = 20; log-rank P = 0.022; univariate Cox HR = 0.459, 95% CI 0.235–0.898, P = 0.023)). (B) Median-dichotomized OS separation (log-rank P = 0.0033). ITGB1 was the sole protein significant in the continuous Cox screen (HR = 0.112, 95% CI 0.043–0.29; BH-FDR = 0.0063 across 958 proteins). (C) OS by pre-treatment ITGB1 within each treatment (FOLFIRINOX log-rank P = 0.0026).

To identify prognostic EV proteins, we screened pre-treatment protein levels against OS using univariate Cox models across 958 proteins. ITGB1 was the only significant protein after BH correction (HR = 0.112, 95% CI 0.043 to 0.290; BH-FDR = 0.0063), with higher pre-treatment ITGB1 associated with longer OS. When using the median as the cutoff, patients with high ITGB1 showed significantly better OS compared to patients with low ITGB1 (log-rank P = 0.0033) (**Fig. 4B**). When the analysis was stratified by treatment, the association was significant within FOLFIRINOX (log-rank P = 0.0026) but not within gemcitabine (**Fig. 4C**), though the smaller gemcitabine subgroup limits this comparison. Notably, ITGB1 was not significantly altered by treatment (post- vs pre-treatment fold change 0.912, adjusted P = 0.268 in the whole cohort; not significant within FOLFIRINOX or gemcitabine), indicating that its prognostic value reflects baseline abundance rather than a treatment-induced change. Because post-treatment biomarkers post-date the time origin, these associations are subject to immortal-time bias and are reported as exploratory.

We next asked whether combining pre-treatment markers improved risk stratification beyond ITGB1 alone. A candidate shortlist of 21 proteins was prepared without reference to survival data. Twenty proteins were drawn from the whole-cohort paired post-versus pre-treatment comparison under relaxed thresholds (BH-FDR < 0.1, |log_2_FC| > log_2_(1.2)) and prioritized by adjusted P value. ITGB1 was added as the twenty-first candidate based on the univariate survival screen described above. A 4-marker risk score was developed, where each marker showed independently associated with OS (**Fig. 4D**). Higher pre-treatment CSPG4 was associated with a higher hazard of death (HR = 4.17, 95% CI 1.61 to 10.76; P = 0.0032), whereas higher ITGB1 (HR = 0.06, 95% CI 0.02 to 0.19; P < 0.0001), RTN4RL2 (HR = 0.28, 95% CI 0.13 to 0.60; P = 0.0012), and TNC (HR = 0.33, 95% CI 0.14 to 0.75; P = 0.0085) were each associated with a lower hazard. Together, these analyses indicate that baseline EV protein abundance, rather than its treatment-induced change, marks a prognostic axis distinct from the platelet response described above.

### A composite platelet signature score quantifies a treatment-specific difference

To quantify the FOLFIRINOX-associated platelet response at the level of individual patients, we summarized the platelet-related proteins identified above into a single platelet signature score, defined as the sample-wise mean z-score of seven proteins that were reduced in the FOLFIRINOX comparison (PF4, PF4V1, PPBP, ITGA2B, ITGB3, GP5, SPARC). Under FOLFIRINOX, the score declined from pre-to post-treatment (Δ = −0.744; paired Wilcoxon P < 0.0001; Cohen’s d ≈ 1.06), whereas under gemcitabine it changed only modestly and non-significantly (Δ = −0.243; P = 0.22; d ≈ 0.32) (**Fig. 5A**). Comparing the change directly between regimens, the decrease was larger under FOLFIRINOX (Wilcoxon rank-sum P = 0.034) (**Fig. 5B**). None of the individual platelet markers differed significantly between treatments, yet the composite score did, indicating that aggregating correlated markers increases sensitivity to this coordinated, treatment-associated change.

**Fig. 5.**
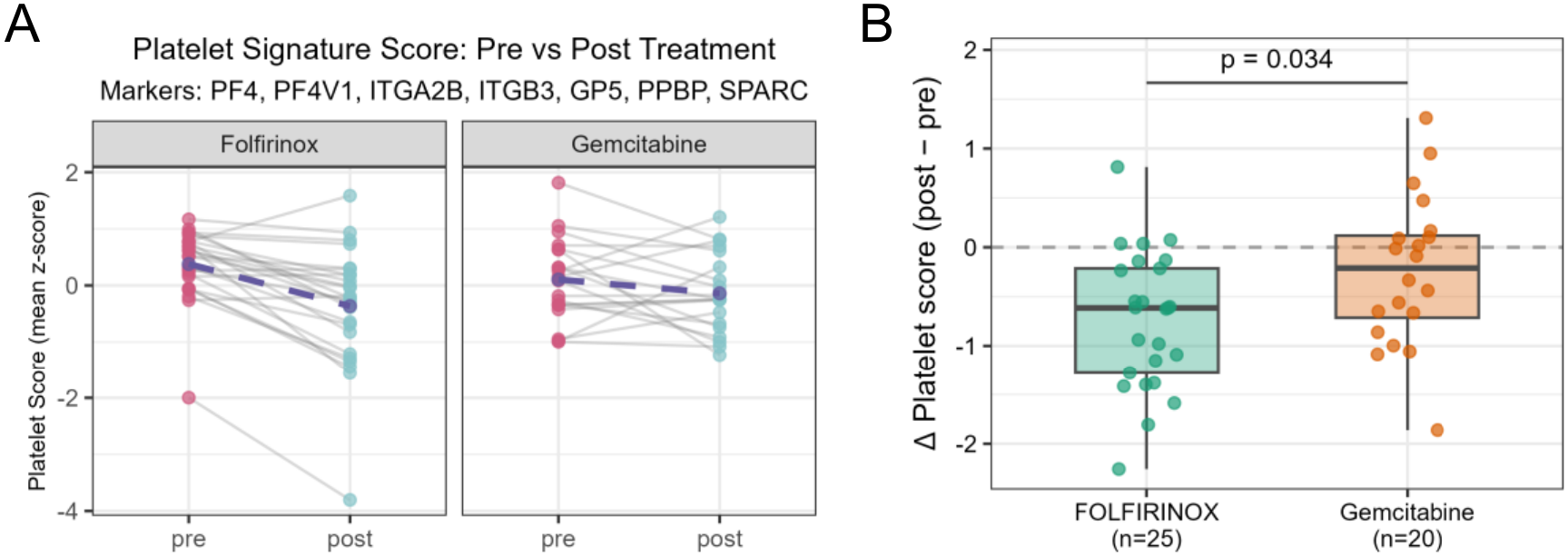
Treatment-specific dynamics of the EV platelet signature score. (A) Patient trajectories of the platelet signature score (sample-wise mean z-score of PF4, PF4V1, PPBP, ITGA2B, ITGB3, GP5, SPARC) from pre-to post-treatment (FOLFIRINOX, n = 25; gemcitabine, n = 20). Grey lines connect paired samples; purple dashed line/point marks the group mean. (B) Between-treatment comparison of the change in platelet signature score (Δplatelet = post − pre). Boxplots show the median and interquartile range with whiskers to 1.5× IQR; overlaid points are individual patients.

### Within-treatment platelet depletion is independently associated with survival

Finally, we asked whether the magnitude of the treatment-induced platelet change was itself prognostic. In a multivariable Cox model (n = 45; 36 events) including age, treatment, and Δplatelet score (post − pre), the Δplatelet score was associated with OS independently of age and treatment (HR = 0.56 per unit; 95% CI 0.335 to 0.945; P = 0.030) (**Fig. 6A**). A higher Δplatelet score, corresponding to less treatment-induced platelet depletion, was associated with lower hazard. Its addition modestly improved the age + treatment model (likelihood-ratio P = 0.028). Bootstrap-corrected discrimination increased slightly (Harrell’s C-index 0.603 to 0.616; 1000 iterations). Age and treatment were retained as adjustment covariates; because treatment assignment was non-random and FOLFIRINOX recipients were younger, the treatment term is not interpreted causally.

**Fig. 6.**
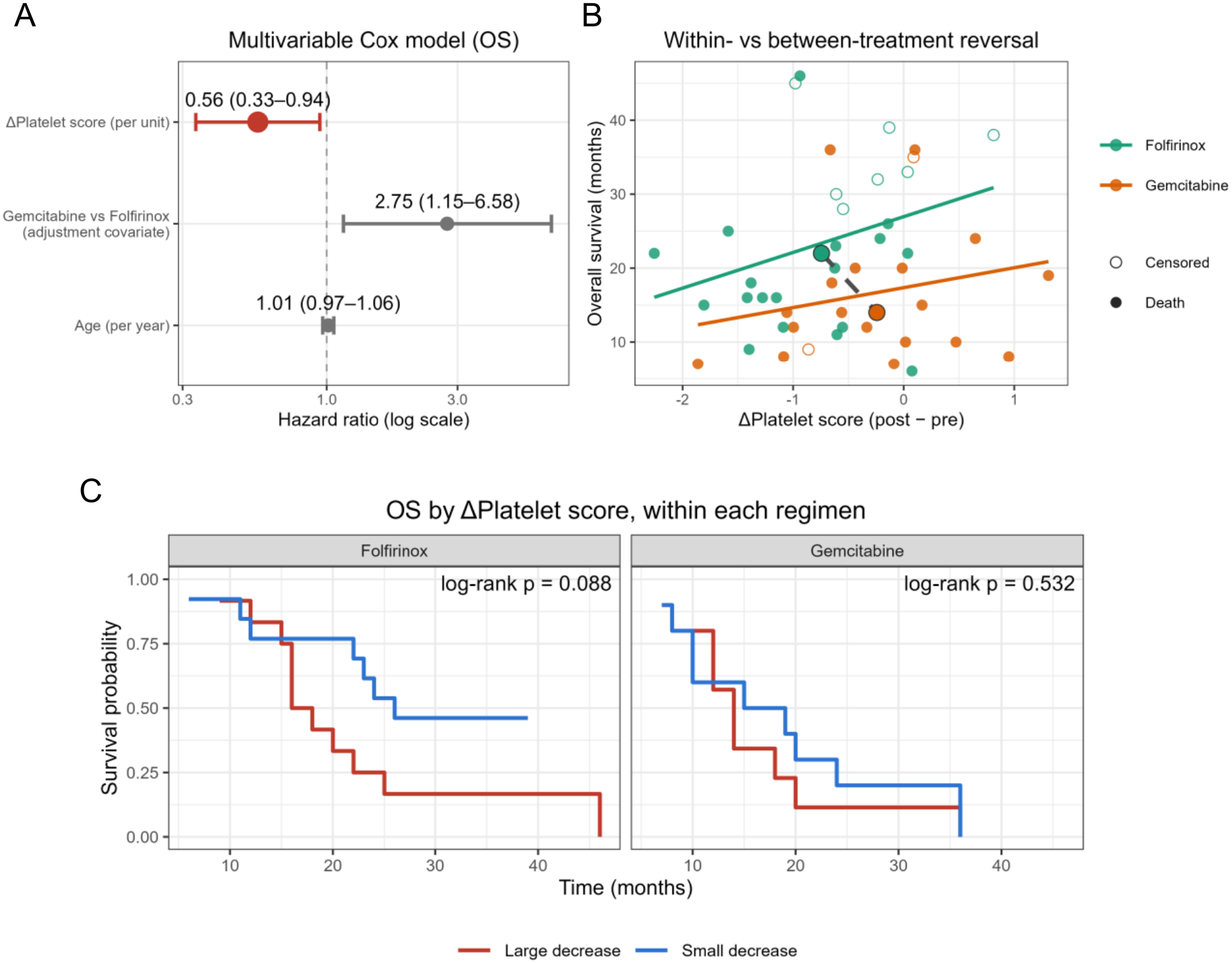
Treatment-induced change in the EV platelet signature carries prognostic information independent of age and treatment. (A) Forest plot of the multivariable Cox model for OS (n = 45; 36 events), including age, treatment, and Δplatelet score (post − pre). Points are hazard ratios (HR) and horizontal bars are 95% confidence intervals (CIs). Age and treatment are included as adjustment covariates. (B) Relationship between Δplatelet score and OS. Solid lines are within-treatment linear trends, and the dashed grey line connects the two treatment centroids (mean Δplatelet score, median OS). (C) Kaplan-Meier OS within each treatment group, with patients dichotomized at the regimen-specific median Δplatelet score (large vs small decrease).

Because the model adjusts for treatment, this association reflects a within-treatment effect. In other words, among patients receiving the same regimen, a more negative Δplatelet score (greater treatment-induced platelet depletion) was associated with shorter survival. This individual-level relationship is distinct from the group-level pattern, in which FOLFIRINOX showed both the largest platelet depletion and longer OS (**Fig. 6B**). Plotting Δplatelet score against OS within each regimen shows a consistent within-treatment trend (less depletion tracking with longer survival), whereas the two treatment centroids run in the opposite direction. The group-level pattern reflects differences in treatment assignment, while the within-treatment pattern isolates patient-to-patient variation once regimen is held constant. Consistent with the adjusted model, dichotomizing patients at the regimen-specific median Δplatelet score reproduced this within-treatment direction in both groups. Within the FOLFIRINOX group, median dichotomization of Δplatelet score showed a concordant but not significant trend (log-rank P = 0.088; n = 25, 18 events) (**Fig. 6C**), and the gemcitabine-based group was underpowered. Together, these results link the FOLFIRINOX-specific platelet EV response to patient outcome. Because the platelet signature was defined from the same cohort, these prognostic associations are exploratory and will require validation in an independent cohort.

## Discussion

Mag-Net enriches EVs by binding negatively charged phospholipids such as phosphatidylserine [30] and can quantify more than 4,000 proteins from fresh plasma samples when coupled with DIA-MS. Compared with differential centrifugation and size-exclusion chromatography, its compatibility with automation and low sample input requirement (10–100 µL) make it attractive for high-throughput plasma proteomics. Recent improvements, including KingFisher-based automation and Evotip loading after digestion, have enhanced scalability and reproducibility, with an average protein-quantification CV of 18% across two 96-well plates and paired-sample correlations ranging from r = 0.857 to 0.991 [24]. We quantified an average of 1,186 proteins per sample, consistent with recent reports [33, 34]. This difference highlights the susceptibility of bead-based plasma and serum proteomics to preanalytical variation, including platelet and PBMC contamination, centrifugation protocols, storage duration, and freeze–thaw cycles [34–37]. Standardized operating procedures are therefore essential to improve the consistency and reproducibility of EV proteomic measurements.

EV-based monitoring may provide a clinically relevant means of tracking molecular responses to pancreatic cancer treatment. Our findings suggest that Mag-Net isolated serum EVs can reveal chemotherapy-associated protein changes and capture treatment-related biological responses. Previous pancreatic cancer studies using ultracentrifugation identified tumor-associated proteins such as GPC1, TIMP1 and THBS2, but largely focused on diagnosis or prognosis rather than treatment monitoring [11]. However, ultracentrifugation may co-isolate non-vesicular proteins and lipoproteins, whereas CD9-based immunoaffinity methods can improve selectivity but capture only specific EV subpopulations [19]. Because ultracentrifugation, size-exclusion chromatography, and immunoaffinity approaches differ in EV recovery, purity, and proteomic coverage [38], standardized workflows such as automated Mag-Net enrichment combined with quantitative proteomics may improve reproducibility and support the clinical validation of EV biomarkers for treatment monitoring.

Using the established Mag-Net coupled with DIA-MS workflow, we have evaluated longitudinal changes of the EV-enriched proteome in paired serum samples of patients with PDAC receiving systemic therapy. Most patients received either FOLFIRINOX or gemcitabine-based therapy except for six patients receiving other regimens. Patients treated with gemcitabine-based therapy were older than those treated with FOLFIRINOX. This difference is consistent with clinical practice, because gemcitabine-based treatment is often selected for patients who may not tolerate intensive combination chemotherapy. Treatment was not randomly assigned, so differences in survival or proteomic changes between treatment groups cannot be interpreted as direct treatment effects. FOLFIRINOX has shown greater efficacy than gemcitabine in selected patients with metastatic PDAC [39], but those randomized trial results do not make the treatment groups in this observational cohort directly comparable.

The paired analysis identified 26 proteins that changed after treatment. Proteins that increased after treatment were mainly related to extracellular matrix and cell adhesion, including LAMA2, LAMC1, FBLN2, SVEP1 and RELN [40–46]. Proteins that decreased included regulators of the actin cytoskeleton, such as PFN1, ANXA7 and SRI [47–49]. Pathway analysis supported these findings: extracellular matrix organization and cell adhesion were increased, whereas cytoskeletal organization, coagulation, and hemostasis were reduced. These changes may reflect remodeling of the tumor microenvironment, changes in stromal or endothelial activity, or systemic effects of chemotherapy. Treatment-associated changes were more pronounced in the FOLFIRINOX group. Thirty-two proteins changed in the paired FOLFIRINOX analysis, whereas no individual proteins reached the same statistical threshold in the gemcitabine analysis. FOLFIRINOX-associated changes included increased extracellular matrix and structural proteins, consistent with the combined analysis.

A notable finding was the coordinated reduction in platelet- and coagulation-associated proteins after FOLFIRINOX. These included PF4, PF4V1, PPBP, GP5, ITGA2B, and ITGB3. Pathway analysis also identified platelet activation, coagulation, and hemostasis as down-regulated processes. The gemcitabine group analysis showed changes involving cytoskeletal, microtubule, and cell-cycle pathways but not the same platelet/coagulation program. Platelets are involved in several aspects of cancer progression [50, 51]. They can interact with circulating tumor cells, support tumor-cell survival, promote vascular adhesion and extravasation, and contribute to metastasis and immune evasion. Platelet-derived EVs may also transport proteins and signaling molecules that affect tumor cells and the tumor microenvironment. The reduction in platelet-associated proteins after FOLFIRINOX could reflect thrombocytopenia, reduced platelet activation, altered release or clearance of platelet-derived EVs, or changes in co-isolated platelet material but these may also occur after gemcitabine-based therapy. The composite platelet score provided a more sensitive measure of this coordinated response than any single protein. The score decreased substantially after FOLFIRINOX but showed a smaller, nonsignificant decrease after gemcitabine-based therapy. Combining correlated proteins may improve the detection of biological processes in a modest-sized cohort. However, because the score was developed and evaluated in the same dataset, its effect size may be overestimated. Validation with targeted proteomics, blood counts, and platelet-function measurements is required.

Baseline EV-protein levels showed a different relationship with outcome than treatment-induced changes. Higher pretreatment ITGB1 (CD29) abundance was associated with longer OS and remained significant after correction for multiple testing. ITGB1 encodes integrin β1, which contributes to cell–matrix adhesion and signaling and can influence tumor-cell survival, migration, invasion, and treatment response [52–55]. The association between ITGB1 and longer survival may reflect differences in tumor–stroma interactions or in the cellular sources of circulating EV-associated proteins. However, this relationship may vary according to tumor type, disease stage, treatment, and cell of origin. ITGB1 was not significantly changed by treatment, supporting its interpretation as a possible baseline prognostic marker rather than a pharmacodynamic marker. An exploratory four-protein risk score included ITGB1, CSPG4, RTN4RL2, and TNC. CSPG4, also known as NG2 proteoglycan, is a cancer-associated proteoglycan involved in tumor-cell growth, migration, invasion, angiogenesis, and extracellular matrix interactions [56–59]. TNC is an extracellular matrix protein associated with tissue remodeling, inflammation, and tumor progression [60–64]. The role of RTN4RL2 in PDAC and EV biology is less established. Because these markers were selected and evaluated in the same cohort, the risk score may be affected by overfitting. The reported hazard ratios should therefore be considered hypothesis-generating and require validation in an independent patient cohort.

### Limitations of This Study

This study has several limitations. First, the cohort was small, particularly for the gemcitabine-based group and the other treatment regimens, and the number of events limited the complexity and stability of multivariable survival models. The relatively small cohort also reflects the limited availability of paired pre- and post-treatment PDAC serum samples from a single medical center, and a small number of patients receiving specific treatment regimens. Second, treatment assignment was not randomized, and the FOLFIRINOX group was younger than the gemcitabine group. Consequently, the observed survival difference between treatments cannot be interpreted causally. Third, there is incomplete information regarding disease stage and other clinical variables, the lack of direct platelet counts and coagulation measurements, and the inability to assign EV proteins to their cellular source. Finally, the prognostic models were internally evaluated in the discovery cohort and were not externally validated. Despite these limitations, the study demonstrates the feasibility of longitudinal serum EV proteome profiling in PDAC to identify treatment-associated protein biomarkers for guiding effective therapy.

### Conclusions

This study demonstrates the feasibility of serum EV proteomics for discovery of treatment-associated protein biomarkers from minimal serum volumes (only 50 µL) using an automated Mag-Net workflow coupled with DIA-MS. In paired PDAC samples, treatment-associated changes were detectable despite the complexity of serum. FOLFIRINOX was associated with a coordinated reduction in platelet- and coagulation-related EV proteins, whereas gemcitabine showed less pronounced changes involving cytoskeletal, microtubule, and cell-cycle pathways. A composite platelet EV signature was associated with OS after adjustment for age and treatment, while higher pretreatment ITGB1 and an exploratory four-protein panel incorporating ITGB1, CSPG4, RTN4RL2, and TNC supported the potential of EV proteins for baseline risk stratification. All these findings suggest that serum EV proteomics can capture complementary information about pretreatment disease risk and treatment-related pharmacodynamic responses. Future studies will validate these findings prospectively in larger and treatment-balanced cohorts, and integrate EV measurements with clinical, hematologic, and imaging data.

## Acknowledgements

This work was supported by NIH R01CA258240 (to T.S. and D.M.L.) and NIH R33CA287139 (to T.S.). In addition, D.M.L. acknowledges support from the Maud T. Lane Professorship. PNNL is a multi-program national laboratory operated by Battelle for the Department of Energy (DOE) under Contract DE-AC05-76RLO 1830. A portion of this research was performed using EMSL, a national scientific user facility sponsored by the DOE’s Office of Biological and Environmental Research and located at PNNL.

## Author contributions

T.S. and D.M.L. conceived and designed the study. S.M., K.A. and T.T.L. performed all proteomic sample preparation. I.K.A. conducted DIA-MS data acquisition. S.M. and T.T.L. performed proteomic data analysis and interpretation. S.M., J.D. and S.L. performed statistical analysis. V.S. and K.C.C. provided clinical protocol support and collected PDAC patient serum samples. S.L., D.M.L. and T.S. provided input on the experimental design, data presentation, and manuscript preparation. S.M., D.M.L. and T.S. wrote the manuscript with input from all other authors.

## Competing interests

All the authors declare no competing interests.

## Notes

### Competing Interest Statement

The authors have declared no competing interest.

